# CSLAN: Cross-Species Latent Alignment Network for Trauma-Related Cell-Type Classification

**DOI:** 10.1101/2025.06.09.655966

**Authors:** Rui Wu, Alan Shi, Yushi Liu, Alexandre Duprey, Ting Chen, Rui Li, Shuang Cao

**Affiliations:** Hill Research, AI Research Team, 307 Waverley Oaks Rd, Waltham, MA 02452, US; Eli Lilly, Global Statistical Science, Lilly Corporate Center, 893 S Delaware St, Indianapolis, IN 46285; University of Cambridge, Cambridge, CB2 1PZ, UK; University of Illinois Urbana-Champaign, Champaign, IL 61801, US; REGENXBIO, Inc., 9804 Medical Center Dr, Rockville, MD 20850, US; Apple, Inc., Apple Park Way, Cupertino, CA 95014, USA

## Abstract

Understanding cellular identity and function across species is fundamental to deciphering conserved biological principles and translating insights from model organisms to human health. However, integrating single-cell transcriptomic data, particularly from resource-limited human studies, with comprehensive datasets from model organisms like mice remains a significant challenge due to evolutionary divergence and technical variability. Here, we introduce the Cross-Species Latent Alignment Network (CSLAN), a transfer learning framework that effectively bridges this gap for robust cell type classification. CSLAN employs a novel strategy involving focused feature selection on source (mouse) data followed by pretraining an encoder-decoder architecture. Crucially, for transfer to human data, only the encoder is fine-tuned while the latent space processor and the decoder, which encapsulate learned, conserved cell type signatures, remains fixed. This targeted adaptation mechanism allows CSLAN to achieve exceptional accuracy (e.g., 96.67% for human immune cells in a trauma context) by efficiently aligning human gene expression profiles to a biologically informed latent space derived from mouse data. Our findings establish a powerful and broadly applicable paradigm for cross-species knowledge transfer, enabling deeper biological insights from limited human scRNA-seq data by leveraging the wealth of information from model systems, with significant implications for translational research and the construction of comprehensive, cross-species cell atlases.

## 1 Introduction

Unraveling the complexities of cellular identity and behavior across diverse biological contexts is a central goal in modern biology, with single-cell RNA sequencing (scRNA-seq) providing unprecedented resolution into these processes [1, 2]. This technology has illuminated cellular heterogeneity and dynamic states in development, health, and disease, offering transformative potential for medicine [3]. However, realizing this potential, particularly for human studies, is often constrained by practical limitations such as sample accessibility, inherent data sparsity, high dimensionality, and confounding technical variability, all of which can obscure biological signals and hinder robust analysis [4, 5].

Leveraging the extensive and well-characterized scRNA-seq data from model organisms, such as mice, through transfer learning presents a powerful strategy to overcome these limitations in human studies [6, 7]. The evolutionary conservation of many fundamental biological processes and cell types between mice and humans provides a strong rationale for such an approach [8, 9]. Specifically, comprehensive mouse atlases can serve as invaluable references, aiding in the identification of rare or poorly characterized human cell populations, increasing statistical power for detecting subtle biological changes, and providing a framework for dissecting conserved versus species-specific molecular programs [10, 11]. Despite this promise, significant challenges persist in effectively transferring knowledge across species. Existing methods often rely on direct ortholog mapping, which can be incomplete or ambiguous, or involve fine-tuning entire pre-trained models, risking the loss of robustly learned, conserved features and increasing susceptibility to overfitting when human data are scarce [12, 13]. Moreover, the computational burden of analyzing genome-wide expression data remains substantial.

Here, we introduce the Cross-Species Latent Alignment Network (CSLAN), a novel transfer learning framework designed to address these critical gaps by enabling highly accurate and efficient cell type classification in human scRNA-seq data through knowledge distillation from mouse models. CSLAN’s innovation lies in a dual strategy: first, it employs L1-regularized Multinomial Logistic Regression (L1-MLR) to identify a compact, highly informative k-gene signature critical for cell type discrimination independently within both source (mouse) and target (human) domains, thereby focusing on a shared, functionally relevant biological subspace. Second, an encoder-decoder architecture, incorporating a latent-space residual network processor, is pre-trained on the k-gene mouse data to learn robust cell type representations. For transfer, the core insight is to freeze the pre-trained decoder and latent processor—which encapsulate these learned, conserved cell type definitions—and fine-tune only the encoder using the k-gene human data. This targeted adaptation allows the human-specific encoder to efficiently map human cellular states into the pre-defined, biologically meaningful latent space, without requiring direct gene-level correspondence between the encoders of the two species at the point of transfer.

Our central hypothesis is that this selective encoder adaptation, operating on an optimized, reduced gene set, can more effectively navigate inter-species differences, preserve foundational biological knowledge learned from the source domain, and achieve superior generalization on limited target data. By focusing the learning challenge for human data primarily on projecting into a well-structured latent manifold rather than re-learning entire classification boundaries from scratch, CSLAN significantly reduces data requirements and computational overhead, making sophisticated cross-species analysis more accessible. We demonstrate the efficacy of CSLAN using challenging trauma response scRNA-seq datasets from mouse models and human patients [14]. Our framework achieves state-of-the-art accuracy in classifying human peripheral blood mononuclear cells, substantially outperforming conventional baselines. This work establishes CSLAN as a powerful and versatile approach that (1) provides a robust method for cross-species knowledge transfer, (2) enhances the interpretation of limited human scRNA-seq data through the lens of comprehensive model organism datasets, and (3) offers a computationally efficient solution for a critical task in single-cell genomics. The principles underlying CSLAN have broad implications for advancing translational research and building integrated, cross-species understandings of cellular biology.

## 2 Literature Review

The advent of single-cell RNA sequencing (scRNA-seq) has spurred the development of numerous machine learning methodologies for tasks such as cell type annotation, trajectory inference, and perturbation modeling. While models ranging from logistic regression and graph neural networks (GNNs) to advanced deep learning architectures like transformers have demonstrated considerable success within individual studies or species [15, 16, 17, 12, 18], transferring this knowledge across different experimental contexts, particularly across species, presents distinct and significant challenges.

### 2.1 Advancements in Intra-Species and Cross-Study Transfer Learning

Within the same species, transfer learning has proven effective for generalizing models across different studies or applying pre-trained representations to new downstream tasks. For instance, scDeepSort, a GNN-based approach, leverages pre-training on extensive datasets to achieve robust cell type annotation across new experiments, outperforming many earlier methods [17]. Similarly, transformer-based models like Geneformer, pre-trained on large corpora of human scRNA-seq data (e.g., Genecorpus-30M), learn powerful gene and cell embeddings that can be fine-tuned for diverse applications, including cell type classification, perturbation response prediction, and gene network inference [12]. These successes highlight the utility of learned representations when domain shifts are relatively constrained, such as those between different studies or tissues within the same species.

### 2.2 Challenges and Strategies in Cross-Species Transfer Learning

Cross-species transfer learning, particularly between model organisms like mice and humans, introduces greater complexity due to evolutionary divergence, differences in gene regulation, and variations in experimental protocols. Despite these hurdles, several approaches have achieved notable success.

#### 2.2.1 Ortholog-Based Transfer

A common strategy for bridging species involves mapping orthologous genes. Early work demonstrated that relatively simple models, such as L1-regularized logistic regression and multi-layer perceptrons (MLPs), pre-trained on mouse bone marrow scRNA-seq data, could achieve high accuracy (e.g., ∼ 83% zero-shot, *>*90% with minimal fine-tuning on 4-8 human samples per cell type) in annotating human bone marrow cells when using a one-to-one ortholog mapping strategy [16]. This study highlighted the conservation of core transcriptional programs defining hematopoietic cell types. More recently, the transformer-based Geneformer architecture was adapted for mice (Mouse-Geneformer), pre-trained on approximately 20 million mouse scRNA-seq profiles. By converting human gene identifiers to their mouse orthologs (and vice-versa), Mouse-Geneformer demonstrated strong fine-tuning performance on human cell type classification tasks (e.g., 87-99% accuracy across thymus, cerebral cortex, and breast tissues), comparable to models trained exclusively on human data [19].

However, ortholog-based mapping presents inherent limitations. Not all genes possess a clear one-to-one ortholog across species, and restricting analyses to only such genes can lead to information loss from non-orthologous or complex-orthology (e.g., one-to-many, many-to-many) genes [19, 16]. This issue is particularly acute for certain biological tasks or tissues where the overlap of one-to-one orthologs might be limited. Methodologies often discard genes without strict one-to-one orthologs [16] or select a single representative in cases of multiple orthologs [19], potentially biasing the analysis.

#### 2.2.2 Ortholog-Free and Semi-Supervised Approaches

To circumvent the limitations of direct ortholog mapping, alternative strategies have been developed. ItClust, for example, employs a semi-supervised deep embedding clustering approach [15]. It first trains a stacked autoencoder on labeled source (e.g., mouse) data for both reconstruction and clustering, then transfers the learned encoder to the target (e.g., human) domain, initializing clustering centroids based on predictions from the source model and fine-tuning through an unsupervised deep embedding clustering algorithm on the unlabeled target data. While ItClust demonstrates promising cross-species performance without relying on orthologs, its accuracy sometimes lags behind ortholog-based methods in direct comparisons for specific tasks. Other approaches focus on aligning latent spaces learned independently from each species, using techniques from unsupervised domain adaptation to find correspondences without explicit gene matching, thereby aiming to capture higher-order structural similarities in gene expression programs [11].

#### 2.2.3 Advanced Deep Learning Architectures for Cross-Species Integration

Beyond direct transfer for classification, more complex deep learning models aim to integrate multi-species data into shared representations for broader biological inquiry. Variational autoencoders (VAEs) like scGen have been used to learn unified latent spaces that can model cellular responses to perturbations across species [20]. Graph-based methods, such as CAME (Cross-species Alignment of Manifold Embeddings), utilize heterogeneous GNNs to jointly embed cells and genes from different species, leveraging even non-one-to-one gene homologies to improve cell identity assignment across greater evolutionary distances [21]. Similarly, gene-centric alignment tools like SAMap [22] explicitly incorporate gene sequence homology to guide the construction of a joint cell-gene graph, facilitating more accurate alignment of cell states across divergent species. These methods often aim for a deeper biological integration rather than solely task-specific transfer.

### 2.3 Positioning the Current Work

The existing literature underscores a persistent trade-off in cross-species transfer learning: ortholog-based methods can be highly effective but are constrained by gene mapping limitations, while ortholog-free methods offer broader applicability but may not always achieve the same level of precision for specific supervised tasks like cell type annotation. Furthermore, while large transformer models like Geneformer show immense promise, their pre-training is computationally intensive, and their optimal transfer strategy often still involves ortholog mapping.

Our proposed framework, Cross-Species Latent Alignment Network (CSLAN), seeks to combine the strengths of focused, supervised learning with a novel approach to circumvent direct ortholog mapping limitations during the encoder adaptation phase. By first pre-training an encoder-decoder model on a source (mouse) dataset (using an optimized set of k informative genes as discussed in the Methods section), we then freeze the biologically informed decoder and latent-space processor. A new encoder is subsequently trained from scratch (or minimally initialized) using the target (human) data, tasked with projecting human gene expression into the latent space understood by the frozen decoder.

This strategy obviates the need for explicit gene-to-gene mapping between the source-trained encoder and the target encoder, focusing instead on learning a human-specific input transformation that aligns with the conserved, pre-learned cell type signatures in the decoder. This approach aims to harness the robust feature extraction capabilities of deep learning while offering a flexible adaptation mechanism for cross-species cell type classification. CSLAN achieves exceptional predictive accuracy, coupled with significant advantages in terms of reduced data requirements and enhanced computational efficiency.

## 3 Methods

### 3.1 Datasets

#### 3.1.1 Source Domain (Mouse)

The mouse scRNA-seq data was obtained from [14]. Briefly, male C57BL/6 mice (8–12 weeks old) were subjected to a validated model of bilateral lower extremity crush injury with bone homogenate injection and controlled hemorrhagic shock. Peripheral blood mononuclear cells (PBMCs) were collected at 3, 6, and 24 hours post-injury, alongside samples from uninjured littermate controls. Single-cell 3’ cDNA libraries were constructed using the 10x Genomics platform (v2 chemistry). After initial processing via the CellRanger pipeline and Seurat (including QC, normalization, batch correction), the dataset comprised 3597 mouse cells, with expression quantified for 12398 genes. The following 8 cell types were identified and used for pre-training: B, Cd4+ T, Cd8+ T, Mono, NK, NK-T, Neutrophils and RBC.

#### 3.1.2 Target Domain (Human)

Human PBMC data were collected from trauma patients within 4 hours, and at 24 and 72 hours post-injury. Age- and sex-matched healthy volunteers served as controls. A mapping between the data group id and data collection time is constructed as: {0: Control Group; 1: 24h; 2: 72h; 3: 4h}. Single-cell cDNA libraries were prepared using 10x Genomics v3 chemistry. Following CellRanger processing and Seurat-based QC and normalization, the human dataset comprised 17038 genes. The target cell types for classification were: B, Cd4+ T, Cd8+ T, Mono, NK, NK-T and RBC. 10 cell entries are randomly selected from each cell type and each group as the finetuning dataset and 5 cell entries are selected similarly for the test set. The PBMC cell types in both human and mouse scRNA-seq datasets were annotated based on surface markers. They were shared by Dr. Timothy R. Billiar.

#### 3.1.3 Data Splitting

For the mouse dataset, cells were randomly split into training (80%), validation (10%), and test (10%) sets. The training set was used for L1-MLR feature selection and for pre-training the CSLAN encoder-decoder. The validation set was used for hyperparameter tuning. The test set was used to evaluate the performance of the pre-trained mouse model. For the human dataset, cells were split into training (66.7%) and testing (33.3%) sets. The training set was used for fine-tuning the CSLAN encoder. The testing dataset was used for final evaluation of the transfer learning performance.

### 3.2 Data Preprocessing

#### 3.2.1 Z-Score Normalization

The feature-wised Z-score normalization method is adopted to remove the scale factors, making the analysis of different input features more consistent and the downstream analyzes more robust. In our dataset, the gene expressions are stored in the matrix format *X* with size *n × f*, where *n* is the total number of cells in the dataset and *f* is the amount of input genetic features. The mean *µ*_*j*_ and standard deviation *σ*_*j*_ of gene *j*, can be calculated based on the training dataset by

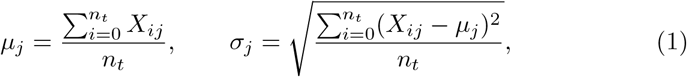

where *n*_*t*_ is the size of the training set and

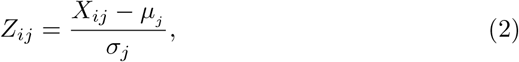

where *Z* is the normalized matrix which helps to facilitate direct comparisons across genes by transforming data onto a common scale.

For L1-MLR feature selection, all 12398 mouse genes were normalized across the training set, and the same process was applied to the human data. For training and inference with the CSLAN encoder-decoder model (operating on k selected genes), the k gene expression values were Z-score normalized similarly. Specifically, for both mouse and human data, means and standard deviations were calculated only from the training set and applied to test and validation dataset.

### 3.3 Informative Gene Selection via L1-Regularized MLR

Feature selection is critical for scRNA-seq data analysis due to the large dimensionality and inherent sparsity of gene expression data. To identify a compact and highly predictive subset of k genes from the mouse dataset, we employed L1-regularized Multinomial Logistic Regression (L1-MLR) to get the most biologically relevant genes. Considering L1 regression’s sensitivity to scale differences between features and in order to encourage fast convergence with the stochastic average gradient descent algorithm, Z-score standardization (as described in section 3.2.1) was applied on the entire dataset for the feature selection step.

We first fitted a PyTorch-based multinomial logistic regression model to the standardized gene expression data. The logistic regression model predicts the probability of each cell belonging to one of the *C* cell types by minimizing the multinomial cross-entropy loss. To enhance the interpretability and sparsity of the model and to effectively identify genes that significantly contribute to distinguishing between cell types, we incorporated an L1 regularization penalty into the logistic regression optimization. Specifically, the L1 penalty term is the sum of the absolute values of the logistic regression coefficients (weights), multiplied by a hyperparameter controlling the strength of regularization. The L1 regularization term encourages sparsity by driving the coefficients (weights) of less informative genes towards zero. The objective function combines the multinomial cross-entropy loss 3 with the L1 penalty on the coefficient vector *β*.

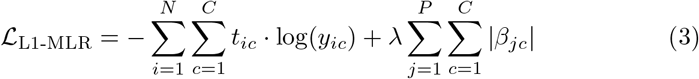

where *N* is the number of cells, *C* is the number of cell types, *P* is the total number of genes, and and *λ* is the L1 regularization strength hyperparameter. *t*_*ic*_ represents the one-hot encoded values of the true cell type label for the *i*^*th*^ cell, with value 1 for the true cell type and 0 values for the others. *y*_*ic*_ is the predicted probability distribution generated by applying the softmax function to the linear logits (*z*_*ic*_):

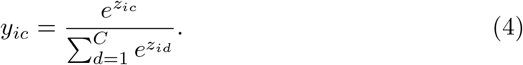

After training the L1-MLR model, the overall significance of each gene *j* was computed by summing the absolute values of its associated regression coefficients across all cell type logits:

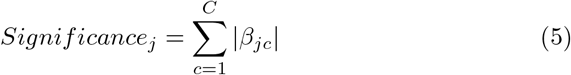

Genes were then ranked by this significance score and the top k genes with the highest scores were selected for downstream analysis. This selection step ensured that only the genes most predictive of cell type identity were retained, significantly reducing the dimensionality of the dataset while preserving essential biological information.

### 3.4 CSLAN Encoder-Decoder Model Architecture

The CSLAN employs an encoder-decoder neural network architecture designed specifically to classify scRNA-seq data into distinct cell types. The encoder module is responsible for transforming the input genetic features, initially represented as high-dimensional gene expression vectors, into a lower-dimensional latent representation. This latent representation captures the most salient biological features and relationships among genes, effectively compressing complex expression patterns into a concise and meaningful manifold.

The decoder module subsequently projects the learned latent representation back into real space to produce cell-type predictions. Specifically, the decoder translates the latent embeddings into probability distributions over the set of possible cell types using feedforward neural networks (FNN) coupled with a softmax activation function.

In FNNs, each such layer processes its input 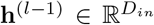 through a sequence of operations. First, a linear transformation is applied:

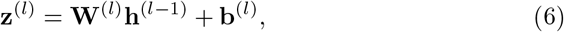

where 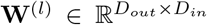 is the weight matrix and 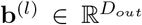 is the bias vector for the *l*^*th*^ layer, yielding a *D*_*out*_-dimensional pre-activation vector **z**^(*l*)^. To stabilize training and improve generalization, batch normalization [23] is subsequently applied to these pre-activations:

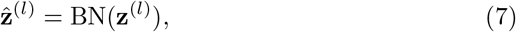

where BN denotes the batch normalization transformation. Following normalization, a Rectified Linear Unit (ReLU) activation function [24] introduces non-linearity:

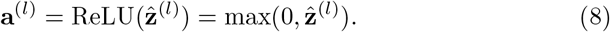

Finally, to mitigate overfitting during the training phase, dropout [25] is applied. A binary mask 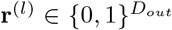 is generated, where each element 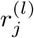 is drawn from a Bernoulli distribution with keep probability *q* (where *q* = 1 − *p*_*drop*_ and *p*_*drop*_ is the dropout rate). The layer’s output, **h**^(*l*)^, is then computed using the inverted dropout scheme:

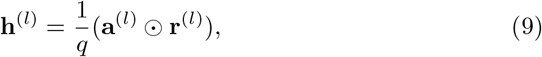

where ⊙ represents element-wise multiplication. During inference, dropout is deactivated, and the output is **h**^(*l*)^ = **a**^(*l*)^, as the scaling by 1*/q* is inherently accounted for by the training procedure. The encoder comprises several such FNN layers, progressively reducing dimensionality from the initial k input features to a 64-dimensional latent space. Similarly, the decoder consists of several FNN layers, which map the processed latent representation towards the output cell type probabilities.

To improve model performance, stability, and convergence speed, the latent representation stage incorporates a residual network architecture [26]. Residual connections enable the model to learn identity mappings by introducing skip connections that allow gradients to propagate more effectively through deeper layers. Formally,

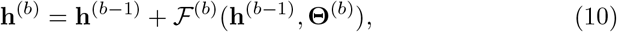

where **h**^(*b*−1)^ and **h**^(*b*)^ represent the input and output feature maps of the *b*^*th*^ residual block, respectively. ℱ^(*b*)^ denotes the residual function parameterized by its set of weights and biases **Θ**^(*b*)^. By integrating residual connections into the latent embedding layers, the model is capable of capturing more nuanced gene interactions while alleviating issues commonly associated with deep neural networks, such as vanishing gradients and diminished training stability.

Overall, the combination of the encoder-decoder structure with residual connections enhances the network’s capacity to model complex biological relationships inherent to single-cell gene expression data, thereby achieving superior predictive accuracy and robustness in cell-type classification tasks.

The overall architecture is illustrated in Figure 1.

**Figure 1.**
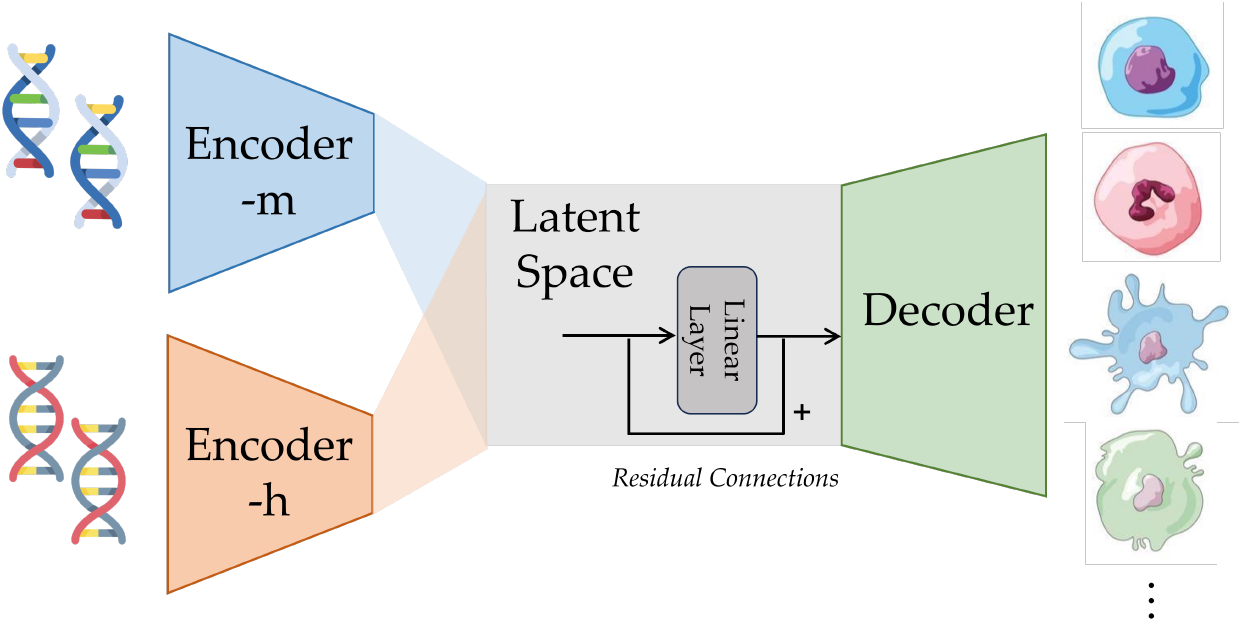
Overall illustration of the classification model, where the Encoder-m and Encoder-h denote the feature embedding layers for mouse and human data respectively.

### 3.5 CSLAN Pre-training on Source (Mouse) Data

At each training step, RNA sequences are randomly selected and fed into the model. To improve performance, individual sequences are weighted in the random sampler to address class imbalance in the dataset, ensuring that minority cell types are adequately represented.

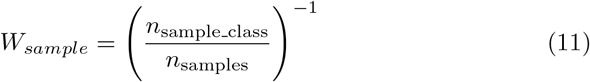

where *W*_sample_ is the individual weight of each sample used in the weighted random sampler, *n*_sample class_ is the number of samples of the same cell type as the current sample, and *n*_samples_ is the total number of samples in the dataset.

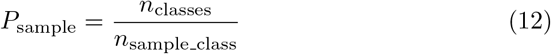

where *P*_*sample*_ is the probability that an individual sample will be chosen by the random sampler, *n*_classes_ is the number of unique cell types in the dataset.

Notably, the use of a weighted sampler implies that one training epoch corresponds to the model processing a number of samples equal to the total size of the dataset, rather than guaranteeing that each individual sample is observed exactly once per epoch. Specifically, samples from underrepresented cell types are presented more frequently, while those from common cell types are shown less often, promoting balanced training across diverse cell populations.

The model’s predictive performance is evaluated by comparing the predicted cell-type probabilities to their corresponding ground-truth labels, which is quantified using the categorical cross-entropy loss function. To minimize the loss and optimize model parameters, we utilized the PyTorch AdamW optimizer, initialized with a learning rate of 5e-4 and a weight decay of 1e-4. A StepLR scheduler is implemented with step size 2.

Following each epoch of training, model performance was assessed on a separate validation dataset using cross-entropy loss. Early stopping was employed based on the cross-entropy loss on the mouse validation set, selecting the model weights from the epoch with the lowest validation loss. This best-performing model was subsequently evaluated on an independent test set to obtain an unbiased estimate of its generalization performance.

Inference of cell types from novel RNA-sequencing data requires the previously selected gene features, along with the statistical parameters (mean and standard deviation) computed from the original training dataset and the optimized model weights.

To perform inference, new RNA-sequencing data are first filtered to retain only those genes identified during the feature selection step. Gene expression values absent from the new data are imputed with zeroes, as the model presumes undetected genes have expression levels below detection thresholds. Subsequently, the filtered expression values are standardized using the mean and standard deviation derived from the original training set. It should be noted that this standardization does not necessarily yield zero mean or unit variance for the inference data. The filtered and standardized data are then passed through the trained model, producing probabilistic predictions for each cell type. The final classification decision corresponds to the cell type assigned the highest predicted probability.

To determine the optimal number of features k, we evaluated the performance of the neural network classifier trained solely on mouse data using varying numbers of top-ranked genes (k=100, 200, 500, 1000). Based on these experiments (Results Section 4.1), k=100 genes were selected, providing the best balance of predictive accuracy and dimensionality reduction. These k genes were used for all subsequent CSLAN pre-training and transfer learning.

### 3.6 Evaluation Metrics

Model performance in cell type classification was rigorously assessed using several standard metrics.

The overall accuracy was calculated as the proportion of correctly classified cells out of the total number of cells. In a multi-class setting with *C* classes, this can be expressed as:

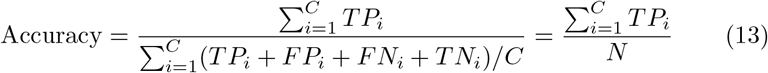

where *TP*_*i*_ is the number of true positives, *FP*_*i*_ is the number of false positives, *TN*_*i*_ is the number of true negatives, *FN*_*i*_ is the number of false negatives for class *i*, and *N* is the total number of samples.

To account for potential class imbalances and provide a balanced measure of performance across all cell types, the macro-average F1 score was calculated. This involves calculating the F1-score for each individual cell type and then averaging these scores. For each class *i*, the Precision (*P*_*i*_) and Recall (*R*_*i*_) are first determined:

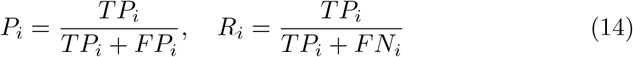

The F1-score for class *i* (*F* 1_*i*_) is then the harmonic mean of its precision and recall:

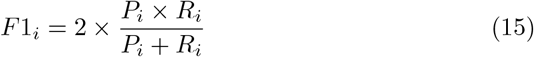

Finally, the F1-Score is calculated as the unweighted mean of the F1-scores for all *C* classes:

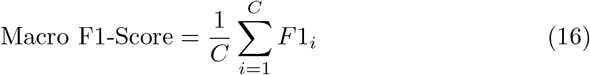

In addition to these summary metrics, confusion matrices were generated to provide a detailed visualization of classification performance for individual cell types, revealing patterns of correct and incorrect predictions between classes.

For latent space and data visualization, Uniform Manifold Approximation and Projection (UMAP) [27] plots were generated to project high-dimensional cell embeddings into a two-dimensional space, facilitating the qualitative assessment of cell type separation and inter-species alignment.

## 4 Results

### 4.1 Optimal Gene Subset Selection for Robust Mouse Cell Type Classification

To establish an effective and computationally efficient input feature set for our Cross-Species Latent Alignment Network, we first optimized the number of genes (k) used for classifying cell types within the source (mouse) domain. An L1regularized Multinomial Logistic Regression model was trained on the mouse training dataset (comprising 12,398 genes) to rank all genes based on their importance in discriminating the eight identified mouse cell types. Subsequently, a neural network classifier, identical in architecture to the CSLAN encoderdecoder (trained end-to-end for this validation), was trained and evaluated on a held-out mouse test set using incrementally larger subsets of these top-ranked genes.

The performance of this classifier across different values of k is presented in Table 1. Notably, utilizing the top k=100 genes yielded the highest test accuracy of 96.11% and a macro F1-score of 0.9585. Performance declined with both fewer genes (e.g., k=10, 90.68% accuracy) and a substantially larger number of genes (e.g., k=1000, 90.54% accuracy), indicating that k=100 provides an optimal balance, capturing essential discriminatory information while minimizing noise and dimensionality.

**Table 1.** Mouse cell type classification accuracy on test set with varying numbers of top k genes selected by L1-MLR.

| Number of Genes (k) | Inerence Accuracy (%) | Macro F1-Score |
| --- | --- | --- |
| 10 | 90.68 | 0.9033 |
| 50 | 94.58 | 0.9454 |
| **100** | **96.11** | **0.9585** |
| 500 | 93.18 | 0.9309 |
| 1000 | 90.54 | 0.9018 |

Qualitative assessment via UMAP visualizations of the mouse data further supported this choice (Figure 3). As k increased from 10 to 100, the separation of cell type clusters in the UMAP embedding became clearer. However, further increasing k to 500 or 1000 did not yield substantially improved cluster distinction and, in some cases, appeared to introduce more diffuse cluster boundaries, consistent with the quantitative performance metrics. Based on these comprehensive evaluations, k=100 genes were selected as the optimal input feature set for all subsequent CSLAN pre-training and transfer learning experiments.

**Figure 2.**
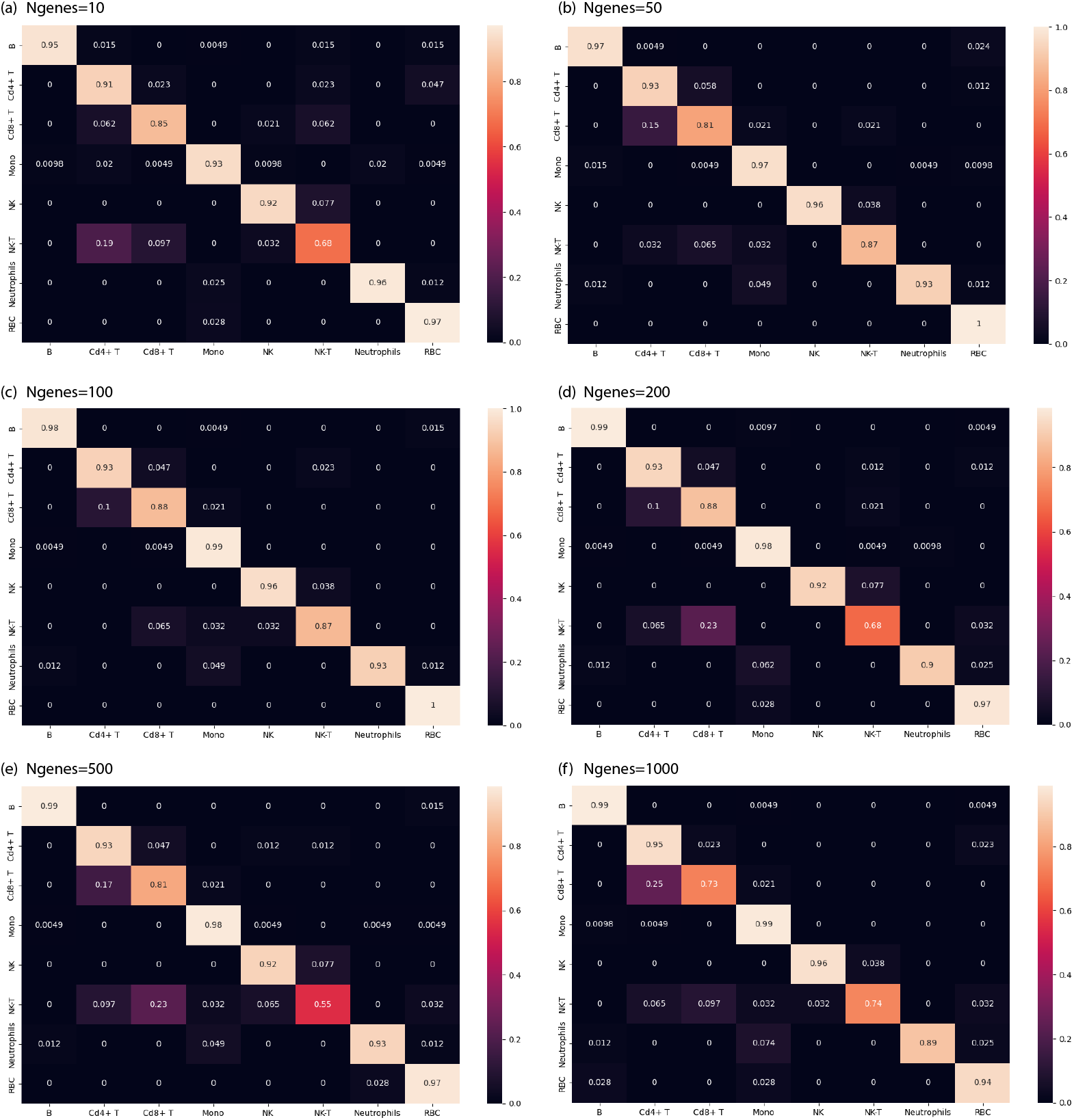
Confusion matrix of the CSLAN model on the held-out mouse test set after pre-training on top-k selected genes, with k varying form 10 to 1000. For each of the confusion matrix, the rows represent true labels and columns represent predicted labels for the 8 mouse cell types. The diagonal’s values denote the accuracy of that specific cell type.

**Figure 3.**
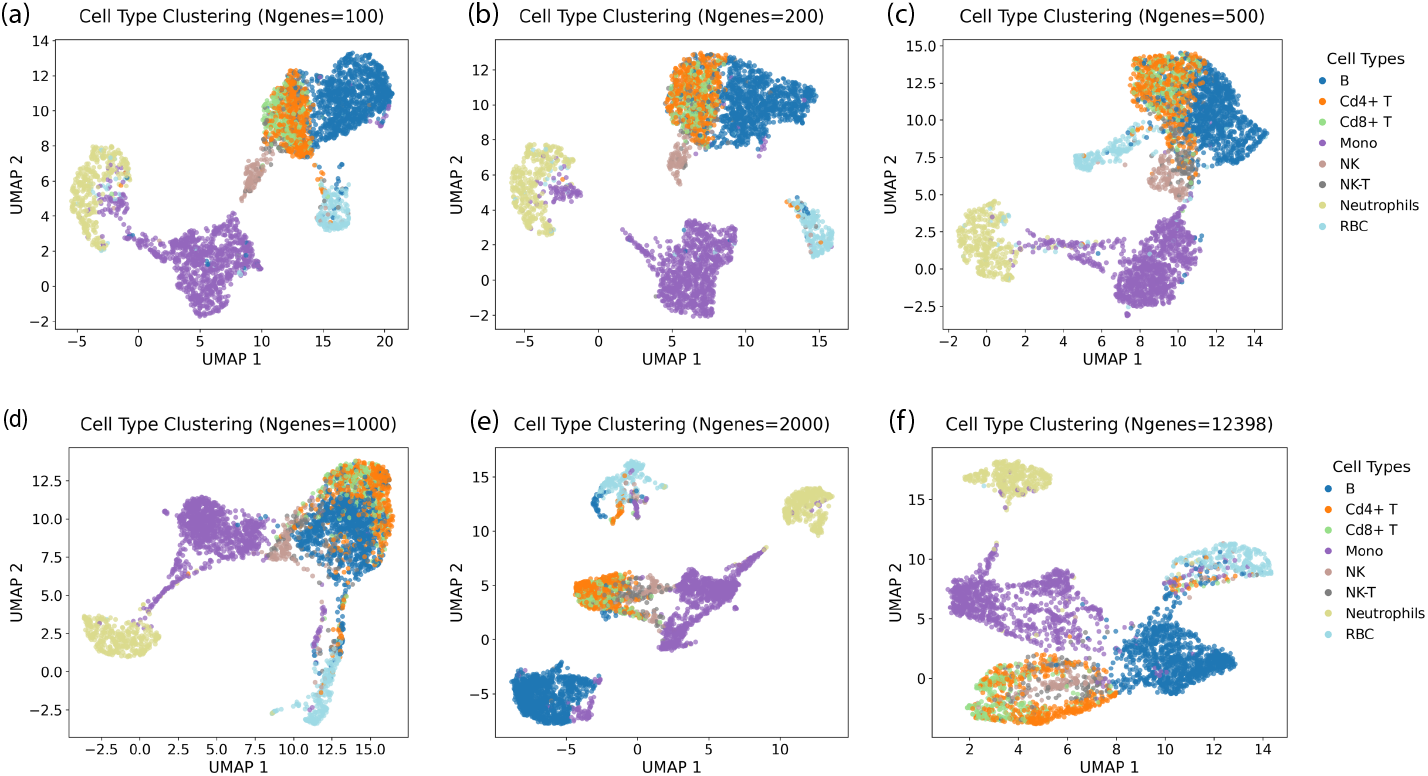
The UMAP of the mouse data with different numbers of top-k genes selected by L1-MLR

### 4.2 Performance of Pre-trained CSLAN on Source (Mouse) Data

Having established the optimal k=100 gene subset, the CSLAN model (encoder, latent RNN processor, and decoder) was pre-trained on the mouse training data. On the independent, held-out mouse test set, the pre-trained CSLAN achieved a robust overall accuracy of 96.11% and a macro F1-score of 0.9585.

The detailed classification performance across the eight mouse cell types is illustrated by the confusion matrix in Figure 2c (corresponding to k=100). High accuracy was observed for most cell types, including B cells (0.98), Monocytes (0.99), NK cells (0.96), Neutrophils (0.93), and RBCs (1.00). Some level of confusion was primarily observed between highly related lymphocyte subsets, such as Cd4+ T cells (0.93 accuracy) and Cd8+ T cells (0.88 accuracy), and between NK cells and NK-T cells (0.87 accuracy). This pattern of minor confusion is consistent with known biological similarities and shared marker expression between these closely related cell populations. Overall, these results confirm that the selected k=100 genes, in conjunction with the CSLAN architecture, are highly effective for accurately classifying mouse cell types in the source domain, providing a strong foundation for subsequent transfer learning.

### 4.3 CSLAN Transfer Learning Performance on Target (Human) Data

The CSLAN transfer learning strategy, involving a frozen decoder and latent RNN with a fine-tuned encoder, was subsequently applied to the human PBMC dataset, using the k=100 human genes. The training and testing curves for this fine-tuning phase are shown in Figure 4. On the human test set, CSLAN achieved an overall accuracy of 96.67% and a macro F1-score of 0.9604 (Table 2).

**Table 2.** Performance comparison on the human PBMC test set (k=100 genes).

| Model | Accuracy (%) | Macro F1-Score |
| --- | --- | --- |
| **CSLAN (Encoder-Tuned)** | **96.67** | **0.9604** |
| Human-Only (100-genes) | 93.33 | 0.9335 |
| Human-Only (all-genes) | 65.83 | 0.6331 |
| Full Fine-tuning (100-genes) | 89.17 | 0.8715 |
| Zero-Shot Transfer (100-genes) | 84.17 | 0.8438 |

**Figure 4.**
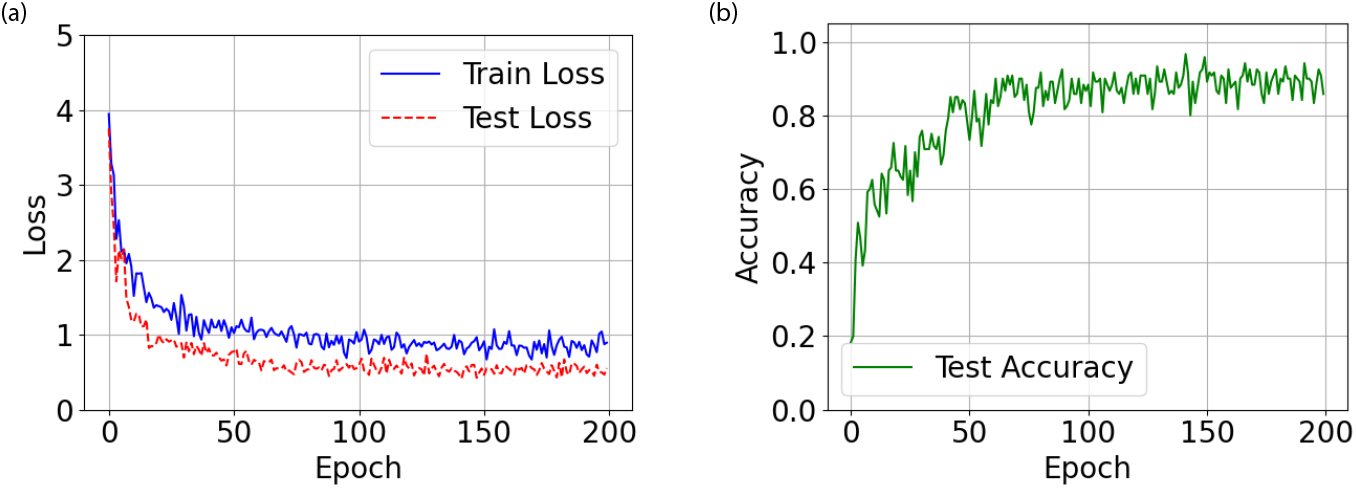
Training and testing loss (left) and accuracy (right) curves for CSLAN during the human fine-tuning phase on the k=100 selected genes. Early stopping was based on validation performance.

CSLAN’s performance was benchmarked against several baseline models, with results summarized in Table 2. Our proposed method significantly outperformed models trained solely on human data. Specifically, training the CSLAN architecture from scratch on only the 100 selected human genes (“Human-Only (100-genes)”) yielded 93.33% accuracy, while training on all available human genes (“Human-Only (all-genes)”) resulted in only 65.83% accuracy, demonstrating that the noise and unsubstantial features will greatly decrease the model’s performance. This latter result underscores the benefit of the feature selection strategy even in a single-species context, in addition to the clear advantage conferred by transfer learning.

Furthermore, CSLAN surpassed alternative transfer learning approaches. A “Zero-Shot Transfer” of the mouse pre-trained model (on 100 genes) to human data achieved 84.17% accuracy, highlighting the necessity of encoder adaptation. “Full Fine-tuning” of all CSLAN components on the 100 human genes yielded 89.17% accuracy. The superior performance of CSLAN (96.67%) compared to full fine-tuning suggests that freezing the decoder and latent RNN effectively regularizes the model, prevents overfitting on the potentially limited human dataset, and preserves robust, conserved biological knowledge learned from the source domain.

CSLAN demonstrates strong performance on the human test set, as evidenced by the confusion matrix (Figure 5). High precision was observed for most cell types, especially for conserved B cells, with slightly lower precision for CD8+ T cells—a finding potentially linked to evolutionary divergence or the subtlety of their k-gene expression signature. Another key outcome of the encoder-only finetuning approach is the achievement of higher and more balanced predictive accuracy across all cell types, compared with the full-parameter finetuning, thereby preventing any single category from exhibiting substantially inferior performance.

**Figure 5.**
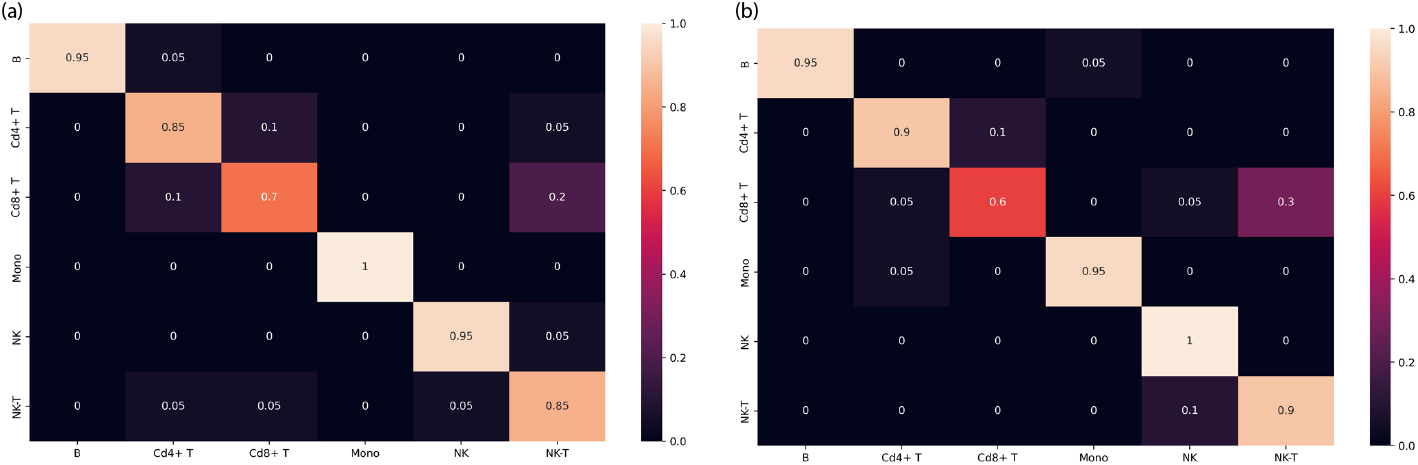
(a) Confusion matrix of the CSLAN model on the human PBMC test set after transfer learning (encoder-tuning) on k=100 selected genes. (b) Confusion matrix of the CSLAN model on the human PBMC test set after full-parameter transfer learning

## 5 Discussion

Our study successfully demonstrated the efficacy of the CSLAN framework, achieving 96.67% accuracy in classifying human cell types by leveraging pretrained mouse models through a targeted encoder-tuning strategy on an optimized gene subset.

A key aspect of CSLAN’s success lies in its ability to bridge the translational gap in leveraging model organism data for human studies, a critical challenge in biomedical research. The robust performance of CSLAN, particularly in classifying human trauma-responsive cells despite known species-specific differences in activation markers, suggests that the framework, by operating on a curated gene set and focusing adaptation on the encoder, captures higher-order functional similarities and conserved core transcriptional programs. This aligns with and extends prior findings that fundamental cell-type-defining gene networks are evolutionarily conserved between mouse and human [9], demonstrating that a strategic latent-space alignment approach can effectively generalize and transfer this conserved knowledge.

The CSLAN architecture and transfer strategy also offer advantages in computational efficiency and model regularization. By freezing the pre-trained decoder and latent-space RNN processor, and fine-tuning only the encoder, CSLAN significantly reduced the number of trainable parameters during the human data adaptation phase. This not only decreased training time compared to end-to-end fine-tuning of the entire network but, more importantly, appeared to mitigate overfitting, as evidenced by CSLAN’s superior performance over the fully fine-tuned baseline, especially when human training data is limited. This targeted tuning preserves the robust, generalizable cell type definitions learned from the larger mouse dataset.

To contextualize the performance of CSLAN, it is instructive to compare its capabilities with established methodologies for cross-species cell type annotation. A notable study by Stumpf et al. [16] demonstrated successful mouse-to-human transfer for bone marrow cell types using L1-regularized logistic regression and multi-layer perceptron models, relying on direct one-to-one ortholog mapping. While acknowledging that direct performance comparisons are nuanced by differences in datasets (trauma-induced PBMCs vs. bone marrow), cell type granularity, and evaluation metrics, their work reported approximately 83% accuracy for zero-shot transfer and over 90% accuracy upon retraining with minimal human data. CSLAN achieved a comparable zero-shot accuracy of 84.17% on our distinct dataset and, critically, attained a significantly higher fine-tuned accuracy of 96.67%. This enhanced performance upon fine-tuning likely reflects the advantages of CSLAN’s deep architecture in capturing complex data structures, the benefits of its initial strategic gene subset selection, and the efficacy of its targeted encoder adaptation mechanism, which is designed to precisely align human cellular states to a robustly pre-defined latent space. Furthermore, CSLAN’s approach, particularly the training of a new human encoder, offers flexibility in scenarios where strict one-to-one ortholog mapping for the entire input feature set might be challenging or suboptimal.

Despite its promising performance, the current study has limitations that open avenues for future research. While CSLAN excelled in classifying major immune cell populations, its efficacy for identifying rare or highly transient cell states, such as specific injury-induced progenitor populations, may be constrained by the inherent scarcity of such cells in training data. It is also worth noting that the reference cell type labels in the trauma dataset were established using graph-based clustering and subsequent marker gene annotation; while a standard approach, this may introduce some degree of label imprecision, particularly for closely related cell types, which could subtly influence model performance metrics like those observed for certain lymphocyte subsets. Future iterations could explore the integration of multi-modal data, such as chromatin accessibility (scATAC-seq) profiles, to provide orthogonal information and potentially improve the resolution of subtle cell states. Furthermore, extending the CSLAN framework to more distantly related non-mammalian species, like zebrafish, would rigorously test its universality in capturing deeper evolutionary conservation of cellular programs.

The broader implications of the CSLAN framework extend beyond the specific application to trauma research. Its demonstrated capacity to decouple species-specific noise from fundamental, biologically meaningful signals offers a valuable blueprint for generalizable single-cell analysis in diverse scenarios where target domain human data is ethically or technically constrained. For instance, CSLAN could potentially accelerate drug discovery and repurposing by enabling the translation of murine drug-response signatures to predict human cellular responses. Moreover, it could contribute to the construction of more comprehensive and integrated cross-species cell atlases, enriching our understanding of comparative cellular biology.

In essence, the CSLAN framework contributes to the ongoing effort to harness the vast knowledge accumulated from model organism research for direct benefit to human health. By refining methods to navigate the complexities of cross-species data integration, we can more effectively translate fundamental biological discoveries into clinical insights.

## 6 Conclusion

This study introduced the Cross-Species Latent Alignment Network (CSLAN), a novel transfer learning framework that effectively leverages knowledge from mouse scRNA-seq data for robust cell type classification in human samples. CSLAN’s strategy, which combines strategic gene feature selection with the pre-training of an encoder-decoder architecture and subsequent targeted finetuning of only the encoder on human data, demonstrated high classification accuracy. This approach successfully bridged the translational gap, outperforming relevant baselines and underscoring the benefits of preserving learned biological knowledge in the decoder and latent processor while adapting the input transformation to the target species.

The efficacy of CSLAN highlights the significant conservation of core transcriptional programs governing cell identity across species and demonstrates a computationally efficient method for managing domain shift and mitigating overfitting, particularly when target human data are limited. The successful alignment of homologous cell types in the learned latent space further validates the capture of species-invariant features critical for cell identity.

While focused on mouse-to-human transfer within the context of immune cell responses to trauma, the principles underpinning CSLAN offer a promising blueprint for broader applications in cross-species single-cell analysis. By facilitating the transfer of fundamental biological insights, CSLAN contributes a valuable tool to the growing arsenal for leveraging model organism data, thereby accelerating discovery and deepening our understanding of human biology in health and disease, and paving the way for advancements in translational research.

## Acknowledgement

The authors thank the ASA-AI/ML Scientific Working Group for suggestion and feedback. The authors thank Dr. Timothy R. Billiar for sharing the scRNA-seq mouse/human data and associated cell types. The authors would like to thank Jason Zhao and Erick Yan for their valuable contributions and support. The authors gratefully acknowledge the support of the Microsoft for Startups program.

